# SdiA as a Repressor of Phagocytosis and Intracellular Survival in *Klebsiella pneumoniae*: Insights from Macrophage and Amoeba Models

**DOI:** 10.64898/2026.05.18.725935

**Authors:** Sergio Silva-Bea, Ricardo Calderon-Gonzalez, Joana Sa-Pessoa, Ana Otero, Manuel Romero, Jose A. Bengoechea

## Abstract

1.

In 2024, the World Health Organisation (WHO) classified *Klebsiella pneumoniae* as a maximum priority pathogen for the development of new alternatives to antibiotics. In this context, understanding the regulation of key virulence mechanisms is essential. Here, we investigated the role of the orphan quorum-sensing receptor SdiA in modulating virulence-associated processes during macrophage infection. Deletion of *sdiA* (Δ*sdiA*) significantly increased susceptibility to phagocytosis, as demonstrated using an amoeba predation model in which mutant strains formed larger clearance zones compared to wild-type bacteria. This phenotype was also observed in murine macrophages, where Δ*sdiA* strains exhibited increased adhesion (1.5 to 2.5-fold) and phagocytic uptake. Reduced uronic acid levels were also quantified in mutant strains, indirectly indicating a diminished capsule production, likely contributing to this enhanced phagocytosis. Despite enhanced uptake, Δ*sdiA* strains showed increased intracellular survival and replication rates within macrophages, leading to reduced host cell viability. This effect occurred despite loss of interbacterial killing capacity against *E. coli*, suggesting that enhanced intracellular fitness is not driven by classical antibacterial offensive mechanisms. Notably, mutant-infected macrophages displayed increased generation of reactive oxygen species (ROS), NF-κB expression, and pro-inflammatory cytokines (mCXCL10 and mTNFα) production, indicating that macrophage defence mechanisms are not impaired during mutant infection. Overall, bacterial survival of Δ*sdiA* could result from overwhelming, rather than actively suppressing, host defences. Together, these findings identify SdiA as a negative regulator of phagocytosis and intracellular survival in *K. pneumoniae* and highlight a context-dependent role in virulence. This work provides new insights into the regulatory networks governing host-pathogen interactions and bacterial adaptation to the intracellular environment.

**Graphical Abstract.**
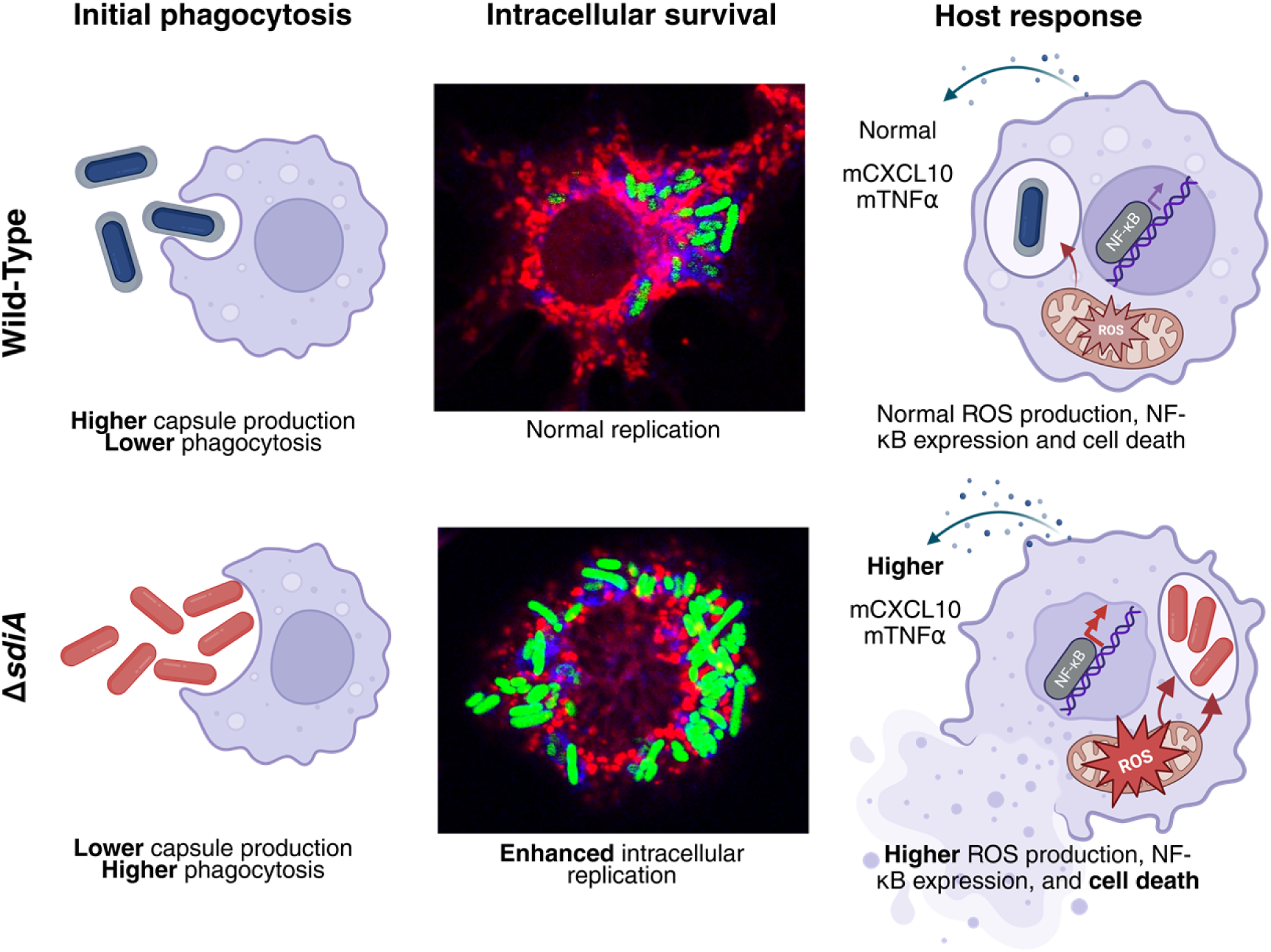
Loss of *sdiA* strongly affects phagocytosis, as mutant strains showed increasing adhesion (1.5 to 2.5-fold) and phagocytic uptake. Diminished capsule production could be contributing to this enhanced phagocytosis, as reduced uronic acid levels were also quantified in mutant strains. Despite being internalized at higher rates, mutants exhibited enhanced intracellular survival and replication, reducing macrophage viability. This fitness advantage occurred independently of classical offensive mechanisms, as evidenced by a lost ability to kill *E. coli*. Notably, mutant-infected macrophages mounted a stronger immune response, marked by elevated ROS, NF-κB expression, and pro-inflammatory cytokines production (mCXCL10 and mTNFα). Together, these findings suggest that strains survive by overwhelming, rather than suppressing, host immune defences. Created with Biorender (https://www.biorender.com/).

**Highlights:**

1. SdiA deletion in *K. pneumoniae* increases susceptibility to phagocytosis.
2. The mutant strains exhibit reduced uronic acid levels, indicative of capsule production.
3. SdiA mutants show enhanced intracellular survival and higher macrophage death.
4. Mutant infected macrophages have higher NF-κB, TNFα, and CXCL10 responses.
5. SdiA-deficient strains lose predatory capacity against *E. coli*.

## 2. Introduction

*Klebsiella pneumoniae* has emerged as a formidable opportunistic pathogen, frequently implicated in a variety of severe hospital- and community-acquired infections (1,2). In 2024, the WHO classified this pathogen as a maximum-priority pathogen due to the increasing prevalence of multidrug-resistant strains, underscoring the urgent need for innovative therapeutic strategies that extend beyond conventional antibiotics and instead target molecular mechanisms involved in its virulence (3–5).

Successful establishment of infection by *K. pneumoniae* relies on its ability to adhere to host cells, evade the immune system, resist phagocytosis, and survive within intracellular environment of immune cells such as macrophages (6). Adhesion and subsequent phagocytic uptake are tightly linked to multiple virulence factors, including surface-associated proteins and, most notably, the polysaccharide capsule, which is known to shield the bacterium from phagocytic uptake by impairing phagocyte attachment, masking surface structures like adhesins, and interfering with complement activation and antibody opsonization, all of which are essential for efficient phagocytosis (3,6,7).

Recent studies have highlighted the importance of the putative Quorum Sensing Acyl-homoserine lactone (AHL) receptor SdiA in regulating adhesion and phagocytosis in *K. pneumoniae* and *E. coli* (5,8,9). Previous work has demonstrated a central role for SdiA in suppressing adhesin expression, including type-1 fimbriae (8,9). Furthermore, SdiA has been shown to promote capsule production in *K. pneumoniae* and *E. coli*, a key phenotype for avoiding opsonisation by antibodies, resisting complement-mediated killing, and evading phagocytosis (3,5,10). Similarly, another study reported reduced adhesion to HeLa cells in an SdiA-deficient *Salmonella enterica* strain (11). Collectively, these SdiA-regulated traits suggest a broad role in suppressing adhesion and phagocytosis, both central processes in immune evasion (3,6,8,9)

Following phagocytosis, the ability of bacteria to survive and potentially replicate within intracellular environments is a key determinant of disease progression. For many years, *K. pneumoniae* was considered strictly an extracellular pathogen, thought to evade phagocytosis primarily through its thick capsule (3,12). However, it is now recognised as a facultative intracellular pathogen capable of actively hijacking macrophages, establishing a specialised niche for replication and eventually spread (6).

*K. pneumoniae* possesses a complex arsenal for survival within the *Klebsiella* Containing Vacuole (KCV). For example, it actively maintains Rab14 on the KCV membrane, a GTPase which marks compartments that are not destined for destruction, avoiding fusion to lysosomes and blocking phagosome maturation into a phagolysosome (12). In addition, the bacterium counteracts host defences through the Type-VI and Type-III Secretion System (T6SS and T3SS) which disrupt mitochondrial network avoiding production of Reactive Oxygen Species (ROS) (13–15). A critical aspect of this host-pathogen interaction is the bacterium’s capacity to modulate the host inflammatory response, for instance through lipid A/LPS modifications or capsule-mediated masking, which reduces Toll-like receptor 4 (TLR4) activation and ultimately limits apoptosis (13,16–18).

Survival alone is insufficient for successful infection; intracellular replication is also required. *K. pneumoniae* manipulates host cell metabolism to enhance nutrient acquisition and support proliferation. It induces increased glycolysis in macrophages (13,19) and upregulates transferrin receptor 1 (TFR1) on the host cell surface, thereby promoting iron uptake, an essential nutrient for bacterial growth and intracellular replication (20). Following replication, *K. pneumoniae* induces necroptosis via the RIPK1/RIPK3/MLKL cascade, resulting in bacterial release and dissemination to new host cells (21). Overall, multiple mechanisms enable *K. pneumoniae* to survive and replicate within macrophages.

Indirect evidence suggests that the Quorum Sensing receptor SdiA may also contribute to intracellular survival and infection progression. In *E. coli*, SdiA can be activated by AHLs, leading to upregulation of the *gad* operon, which enhances acid resistance, an important adaptation for survival in acidic environments such as the gastrointestinal tract or phagolysosomes (8,22). Moreover, SdiA has been implicated in protection against ROS in *E. coli* through enhancing UvrY expression (8). Additionally, AHL-activated SdiA promotes cAMP production via *ydiV* expression, which encodes a protein involved in intracellular signalling (23). cAMP, in turn, contributes to oxidative stress responses by inducing antioxidant enzymes such as catalase and superoxide dismutase (24). In *Enterobacter cloacae*, AHL-activated SdiA has been reported to upregulate a hypothetical protein within a T6SS, although this effect was strongly dependent on culture conditions (25). Similarly, the T6SS effector VasK was downregulated in an SdiA-deficient *Cronobacter sakazakii* strain (26). Furthermore, our previous research demonstrated depletion of virulence of a Δ*sdiA K. pneumoniae* strain in the *Galleria mellonella* infection model in the presence of C6-HSL (5). Collectively, these findings indirectly suggest a potential role for SdiA in promoting intracellular survival and infection progression. However, other studies support the opposite hypothesis, suggesting that SdiA may inhibit intracellular survival and thus act as a negative regulator of infection. For instance, several reports have shown that T3SS and associated effectors in *E. coli* and *Salmonella* spp. are downregulated by SdiA (27–29), while T3SS activity in *S. enterica* is required for intracellular proliferation (30). To date, no studies have examined the impact of SdiA on the intracellular survival of this outstanding pathogen *K. pneumoniae*.

The aim of this study is to advance our understanding of the multifaceted role of SdiA in *K. pneumoniae* pathogenesis, with particular emphasis on its contribution to key infection-related processes including adhesion to phagocytes, phagocytosis, and intracellular survival. To this end, we used a CRISPR-Cas9-generated *sdiA* deletion mutant in the KLEB-33 strain, a multidrug-resistant, highly biofilm-forming, hypermucoviscous strain carrying hypervirulence-associated genes. For comparative purposes, we used the ATCC 13883^T^ strain, which lacks these phenotypic traits. Our findings show that deletion of *sdiA* increases phagocytosis in both amoeba predation model and murine macrophage-infection model. Increased intracellular survival and replication rates in the mutant strain were also observed in macrophages. These results suggest that SdiA acts as a repressor of these virulence-associated phenotypes, potentially through modulation of capsule production, which normally limits host-pathogen interactions. Despite enhanced intracellular survival, Δ*sdiA* strains did not show increased predation of *E. coli* nor impaired macrophage defence mechanisms (ROS and cytokine production). This suggests that the observed intracellular effects may be driven by the higher initial bacterial uptake, which could overcome host capacity and ultimately reduce macrophage viability at late infection stages. Collectively, these findings improve our understanding of the regulatory networks governing *K. pneumoniae* virulence and highlight the complex interplay between bacterial quorum sensing systems and host-pathogen interactions. This knowledge may contribute to the development of anti-virulence strategies targeting bacterial communication and defence mechanisms.

## 3. Materials and Methods

### 3.1. Bacterial strains and culture conditions

In this study, we used the KLEB-33 and ATCC 13883^T^ *K. pneumoniae* strains. KLEB- 33 is a hypermucoviscous, multi-dug resistant (MDR), high biofilm-forming strain that harbours hypervirulence genes (31). ATCC 13883^T^ is a low-biofilm forming, non-MDR strain used for comparative purposes. *sdiA-deficient* strains (Δ*sdiA*) were constructed using a CRISPR-based mutation system (5). Complementation of *sdiA* was performed using a plasmid-based system (Δ*sdiA* + p*sdiA*). *sdiA* was cloned under control of its own native promoter in the low-copy plasmid pME6000 harbouring a tetracycline resistance gene (20 µg/mL). *sdiA* promoter was identified upstream of the gene using the bioinformatic tool BPROM (http://softberry.com). Bacterial strains were routinely grown in Lysogeny Broth (LB) agar (1.5 %) at 37 °C/24 h. Experiment inoculums were grown at 200 rpm for 16 h/37 °C in LB. For each experiment, *K. pneumoniae* strains were re-grown for 2 h in 5 mL LB until mid-exponential phase at 37 °C/200 rpm, centrifuged at 3600 g/20 min, washed with PBS and adjusted to Abs_600 nm_ 1 (**≈** 1.5x10^8^ CFUs/mL). In all the experiments performed the same amount of solvent was added to negative controls, if needed.

### 3.2. *Acanthamoeba* sp. predation assay

Susceptibility to amoeba predation and phagocytosis of bacteria was evaluated as described in the literature with modifications (32). The free-living amoeba *Acanthamoeba* sp. was grown in PYG medium supplemented with kanamycin (60 µg/mL) at 80 rpm, 30 °C/7 days. PYG is composed of 2 % peptone, 0.1 % yeast extract, 0.1 M D-glucose, 0.05 M CaCl_2_, 0.4 M MgSO_4_, 0.25 M Na_2_HPO_4_, 0.25 M KH_2_PO_4_, 0.1 % Na-Citrate, and 0.005 M Fe(NH_4_)_2_(SO_4_)_2_ (ATCC Medium 712). After growth, amoeba was centrifuged (200 g/10 min), washed twice with PBS, and stored at 4 °C. Bacterial inoculum was adjusted to Abs_600 nm_ 1, and 100 µL (**≈** 1.5x10^7^ CFUs) were seeded by extension in agar plates (1.5 %) without nutrients to avoid bacterial growth. A spot of 10 µL of amoeba (≈ 2x10^4^ cells) was placed in the centre of the bacterial-inoculated agar plates and incubated at 30 °C/3 days aerobically. Trophozoite front diameter was measured every day. Morphology of amoebas was checked in the microscope at 72 h incubation at 400x using a coverslip placed over the agar (Dalmau plate technique). The experiment was repeated three times with N=2.

### 3.3. iBMDMs infection assays

Macrophage infection for adhesion, phagocytosis, and intracellular survival dynamics was performed as described in the bibliography (16,33). Immortalised Bone Marrow Derived Macrophages (iBMDMs) cell line derived from wild-type NR9456 mice (BEI Resources, NIAID, NIH) was grown routinely in 25 mL DMEM (Gibco, ref.: 41965) supplemented with 10 % heat-inactivated fetal bovine serum (biowest, ref.: S181B-050) in 80 cm^2^ Tissue Culture treated Flasks (ThermoFisher, ref.: 178905) at 37 °C in a humidified 5 % CO_2_ incubator. Macrophages were routinely tested for *Mycoplasma* contamination. When reached 80 % confluency (**≈** 48 h), cells were washed with PBS, trypsinized with 1 mL 0.25 % Trypsin-EDTA (Gibco, ref.: 25200), counted in Neubauer chamber, and dispensed in 12-well tissue culture plate (Corning, ref.: CLS3512) 5x10^5^ cells/well in 1 mL/well for overnight incubation.

Once macrophages were attached to the bottom of the wells overnight, cells were washed with PBS and infected with bacteria at MOI 50. To synchronise infection, infected plates were centrifuged 5 min/200 g. After 30 min of incubation, infected macrophages were washed with PBS and DMEM supplemented with gentamycin (100 µg/mL) was added to culture to kill extracellular bacteria, and incubated for 60, 180, and 300 min since beginning of infection. 30- and 60-min time-points corresponds to adhered and phagocyted bacteria, respectively. 60-, 180- and 300-min time-point corresponds to intracellular survival dynamics. At each time-point, infected cells were washed with PBS and incubated 5 min with 300 µL of 0.05 % saponin (ref.: S7900-100G) and seeded in LB agar (1.5 %) plates for CFU counting. The experiment was repeated three times with N=2.

### 3.4. Uronic acid quantification

Uronic acid quantification was performed as described in the bibliography (34,35). Briefly, bacterial pellet was lysed with zwittergent 1 % (in 100 mM citric acid pH 2.0) at 50 °C/20 min. Bacterial cellular debris was removed by centrifugation (3600 g/20 min) and supernatant was incubated with 100 % ethanol (1/4 dilution) at -20 °C/20 min to precipitate CPS. CPS was collected by centrifugation (4 °C, 10.000 g/10 min) and dried at 90 °C/5 min. Finally, 1.2 mL of 12.5 mM sodium tetraborate (in pure sulphuric acid) was added to the tubes to dissolve CPS and incubated 100 °C/10 min and cooled to room temperature (RT). To quantify, 20 µl of 0.125 % of Carbazole (in 100 % ethanol freshly prepared) was added and incubated at 100 °C/10 min, mixing every 3 min. Once cooled Abs_530 nm_ was measured. A standard curve of D-Gluconic acid lactone (100 mg/mL serially diluted 1/2 x7 times) was used to quantify the amount of CPS from absorbance data. Inoculums used were seeded in LB agar (1.5 %) to Colony Forming Units (CFUs) count after incubation 24 h/37 °C. Previous experiments indicated no differences in bacterial growth curves between strains and conditions used (5). Micrograms of CPS were expressed per 10^6^ CFU. The experiment was repeated twice with N=3.

### 3.5. Confocal microscopy

For confocal visualisation, iBMDMs were infected with KLEB-33 WT and Δ*sdiA*, performed exactly as described in section **3.3** but with slight modifications, following the experiments described in the literature (12,14). All the steps were performed in dark conditions to avoid degradation of fluorophores. The day before infection macrophages were seeded in 24-well plates (Corning, ref: CLS3526) at 2x10^5^ cells/well (500 µL/well) placing at the very bottom of the well a glass round sterile coverslip. Before infection, the macrophages were washed with PBS and incubated 30 min at 37 °C/ 5 % CO_2_ with MitoTracker (Invitrogen, ref: M7512) following manufacturer recommendations. Macrophages were infected with bacteria (MOI of 50) harbouring plasmid pFPV25.1 for constitutive expression of green fluorescent protein (GFP), with chloramphenicol resistance gene cassette (30 µg/mL) (12). At each time point of infection (60, 180 and 300 min), cells were washed with PBS and fixed with paraformaldehyde (4 %) for 20 min (RT). After fixation, cells were washed with PBS and conserved at 4 °C with NH_4_Cl (14 mM) overnight. Next day cells were washed with PBS and permeabilised with saponin (0.5 % in PBS) for 30 min (RT). For staining, samples were incubated 20 min (RT) firstly with primary antibody Anti-Lamp1 (lysosome-associated membrane protein 1) rat monoclonal IgG_2a_ (1D4B) (Santa Cruz Biotechnology, ref: sc-19992), washed with PBS, and secondly with detection antibody (donkey pAb anti-rat IgG) conjugated to AlexaFluor405 (abcam, ref: ab175670). Both antibodies were used at 0.5 µg/mL diluted in saponin (0.5 % in PBS) with 10 % horse serum. After staining, samples were washed with PBS and incubated overnight in humidity chamber with ProLong™ Gold Antifade Mounting reagent (ref.: P36930) on microscope slides. Finally, samples were sealed with transparent nail polish. Repeatability of experiments was checked with CFU quantification.

A total of 2 replicates per sample were analysed, and 5 pictures/sample were taken in Leica Stellaris 5 at 1000x magnification (0.75x zoom). Analysis of pictures was performed with Leica Application Suite Office (v1.4.6.28433) and Fiji software (v1.54p). Each picture was analysed for quantification of the percentage of infected cells per field, and the number of bacteria counted per infected cell. As GFP signal was recorded outside the macrophages, bacteria were only considered inside the cell if GFP signal colocalised with Lamp1 signal (indicating that bacteria were indeed inside the phagosome) and mitochondria signal surrounded the phagosome.

### 3.6. Macrophage viability

At 300-min time-point of macrophage infection assays (section **3.3**), the cell viability was quantified as described in the literature (16) by incubation for 2 h with Neutral Red colorant (Sigma, ref.: N4638-1G) (40 µg/ml), washed with PBS, and Abs_540 nm_ measurement after solubilization of colorant with destaining solution (1 % glacial acetic acid, and 50 % ethanol). Experiment was repeated three times with N=2.

### 3.7. *E. coli* killing assay

Competition experiments were performed as described in the literature with modifications (15). *K. pneumoniae* (predator) was incubated with *E. coli* (prey) at 10:1 ratio in a dried spot (100 µL) over a pre-warmed agar plate (1.5 %) without nutrients for 5 h/37 °C. Nutrient-free conditions were used to avoid differential growth of bacterial populations and to ensure competition-driven interactions (15). After incubation prey was recovered on LB+ streptomycin (100 µg/mL) and colony counted (CFUs/mL). Streptomycin resistance in *E. coli* MG1655 was induced by repeated exposure to the antibiotic prior to experiments. Sensitivity of *K. pneumoniae* strains to antibiotic and the concentration used was checked before experiments. The experiment was repeated 4 times with N=3.

### 3.8. ROS quantification

Total and mitochondrial ROS quantification was performed in iBMDMs infected with *K. pneumoniae* as described in section **3.3** with modifications. The day before infection macrophages were seeded at 3x10^5^ cells/mL, 200 µL/well in 96-well black clear-bottom plates (Greiner, ref.: 655090). For mitochondrial ROS quantification (14), before infection macrophages were washed with PBS and incubated with 5 µM MitoSOX red (Invitrogen, ref.: 17847181) for 30 min/37 °C. After infection, cells were washed and culture medium was replaced with transparent HBSS (Hanks’ Balanced Salt Solution) (Gibco, ref: 14025092) supplemented with gentamycin (100 µg/mL) to avoid background autofluorescence of DMEM. For total ROS quantification, when gentamycin was added, 5 µM of H_2_DCF-DA (2′,7′-dichlorodihydrofluorescein diacetate) was also added (36). Infected plates were incubated for 300 min at 37 °C/5 % CO_2_ and fluorescence intensity was measured at excitation/emission of 396 nm/610 nm for mitoSOX, and 485 nm/520 nm for total ROS. Repeatability of experiments was checked with CFU quantification. The experiments were repeated 4 times with N=4.

### 3.9. Cytokine quantification

At 300-min time-point of macrophage infection assays (section **3.3**), supernatant was recovered for mCXCL10, mTNFα, and mIL-1β quantification (Preprotech ELISA development Kit, ref.: 900-K153, 900-K54, and 900-K47 for each cytokine, respectively) using TMB Solution I (Biopanda, ref.: TMB-S-001) for colorimetric development. Quantification was performed for each cytokine with a standard dilution curve for each experiment. Experiments were repeated three times with N=2.

### 3.10. THP1-Dual reporter cells

THP1-Dual reporter cells (InvivoGen, cat. no. thpd-nfis) were cultured in Roswell Park Memorial Institute (RPMI) 1640 Medium (Gibco 21875) supplemented with 10 % heat-inactivated fetal calf serum, 10 mM HEPES (Sigma), 100 U/ml penicillin, and 0.1 mg/ml streptomycin (Gibco). Cells were maintained at 37 °C in a humidified 5 % CO₂ incubator. To ensure selection pressure, the growth medium was supplemented with 10 μg/ml of Blasticidin and 100 μg/ml of Zeocin every other passage. For infection experiments, THP1-Dual cells were seeded in 96-well plates at a density of 10^5^ cells/well (200 µL/well) and differentiated into macrophages using 25 ng/ml PMA (phorbol 12-myristate 13-acetate). After 24 hours, the cells were washed with PBS and medium was replaced with antibiotic-free RPMI-1640. After 24 h, cells were infected with *K. pneumoniae* exactly as described in section **3.3**. At 300-min time-point since start of infection, supernatants were harvested to analyse secreted embryonic alkaline phosphatase (SEAP) activity (indicative of NF-κB activation) and luciferase activity (indicative of IFNAR signalling) using QUANTI-Blue^TM^ and QUANTI-Luc^TM^ 4 Lucia/Gaussia reagents, respectively, following manufacturer recommendations (InvivoGen, ref: thpd-nfis). Experiments were repeated three times with N=2. Repeatability of experiments was checked with CFU quantification.

## 4. Results

### 4.1. Δ*sdiA* strains exhibit increased susceptibility to amoeba and macrophage phagocytosis

As an initial approach to investigate the role of *sdiA* in modulating interactions of *K. pneumoniae* with phagocytic cells, we employed a well-established amoeba predation assay using the free-living amoeba *Acanthamoeba* sp. (32). In this model, amoebae plated on a bacterial lawn generate expanding trophozoite halos, the size of which is inversely proportional to bacterial resistance to phagocytosis: resistant strains produce smaller or no halos, whereas susceptible strains generate larger clearance zones (37,38). Consequently, this assay provides a quantitative and visual readout of susceptibility to phagocytic clearance and, by extension, virulence potential (3).

Consistent with the proposed role of SdiA in protecting from phagocytosis as mentioned above (3,8), the Δ*sdiA* strain of ATCC 13883^T^ and KLEB-33 exhibited significantly increased sensitivity to amoeba predation compared to their WT counterparts. This was evidenced by the formation of bigger phagocytic halos in Δ*sdiA* lawns (**Figure 1A**). Complementation of the mutant restored the WT phenotype, indicating a specific SdiA-dependent effect. Microscopic analysis further supported these findings. In WT strains, amoebae were unable to fully clear the bacterial monolayer, with numerous bacterial cells remaining in areas of trophozoite proliferation (**Figure 1B**). These observations suggest that WT strains of *K. pneumoniae* display enhanced resistance to phagocytic predation. In contrast, Δ*sdiA* strains showed near-complete bacterial clearance within the phagocytic halo, indicating reduced resistance to amoeba-mediated predation (**Figure 1B**).

**Figure 1.**
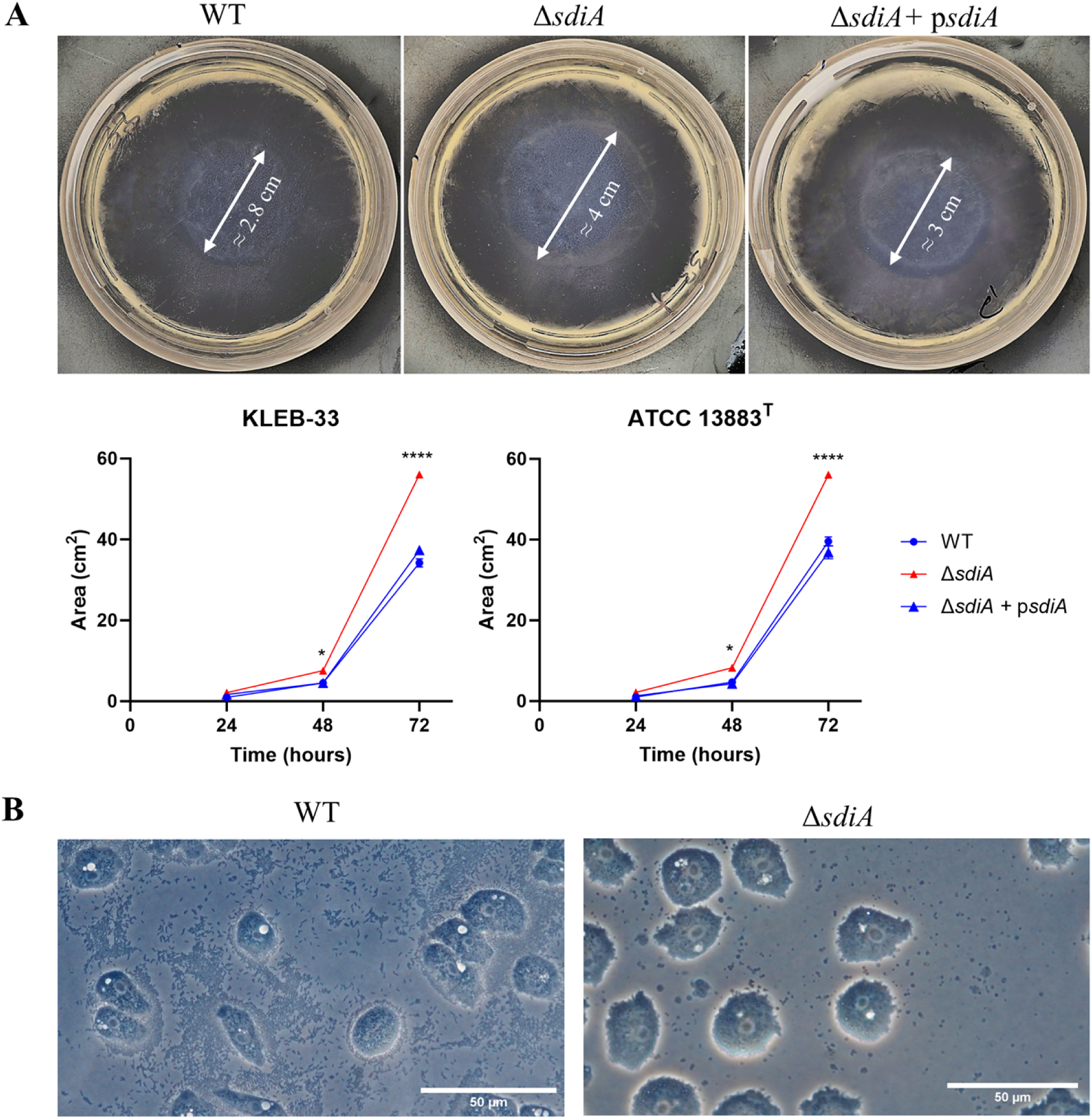
(**A**) Representative images (top) and quantification of halo area (bottom) of predatory halos of amoeba phagocyting WT, Δ*sdiA*, and complemented Δ*sdiA* (Δ*sdiA* + p*sdiA*) of KLEB-33 and ATCC 13883^T^ strains. Images were taken after 48 h of co-culture, illustrating zones of bacterial clearance (“predation halos”) formed by *Acanthamoeba* sp. trophozoites. Halo diameters were measured and expressed as area (cm^2^) to assess susceptibility to amoeba predation. Experiments were performed in three independent assays with N=2 biological replicates. (**B**) Representative images of *Acanthamoeba* sp. predation on *K. pneumoniae* WT and Δ*sdiA* strains. Images were captured at 400x magnification using the Dalmau technique and correspond to predation assays performed with the KLEB-33 strain.

Due to the evolutionary conservation of phagocytosis and intracellular killing mechanisms (37), we next evaluated whether this phenotype was recapitulated in murine macrophages. iBMDMs were infected with WT, Δ*sdiA*, and complemented strains of KLEB-33 and ATCC 13883^T^. Deletion of *sdiA* resulted in a marked increase in macrophage adhesion at 30 min post-infection in both strain backgrounds (**Figure 2**). This was followed by a corresponding increase in phagocytic uptake at 60 min, with Δ*sdiA* strains showing a 1.5- to 2.5-fold higher number of intracellular bacteria compared to WT and complemented strains (**Figure 2**). This trend was consistent across both genetic backgrounds, although the magnitude of the effect varied between strains. In all cases, complementation restored adhesion and phagocytosis levels to those observed in WT infections, confirming that the phenotype is specifically associated with the absence of SdiA. Overall, these data indicate that loss of SdiA enhances both bacterial attachment to macrophages and subsequent internalisation.

**Figure 2.**
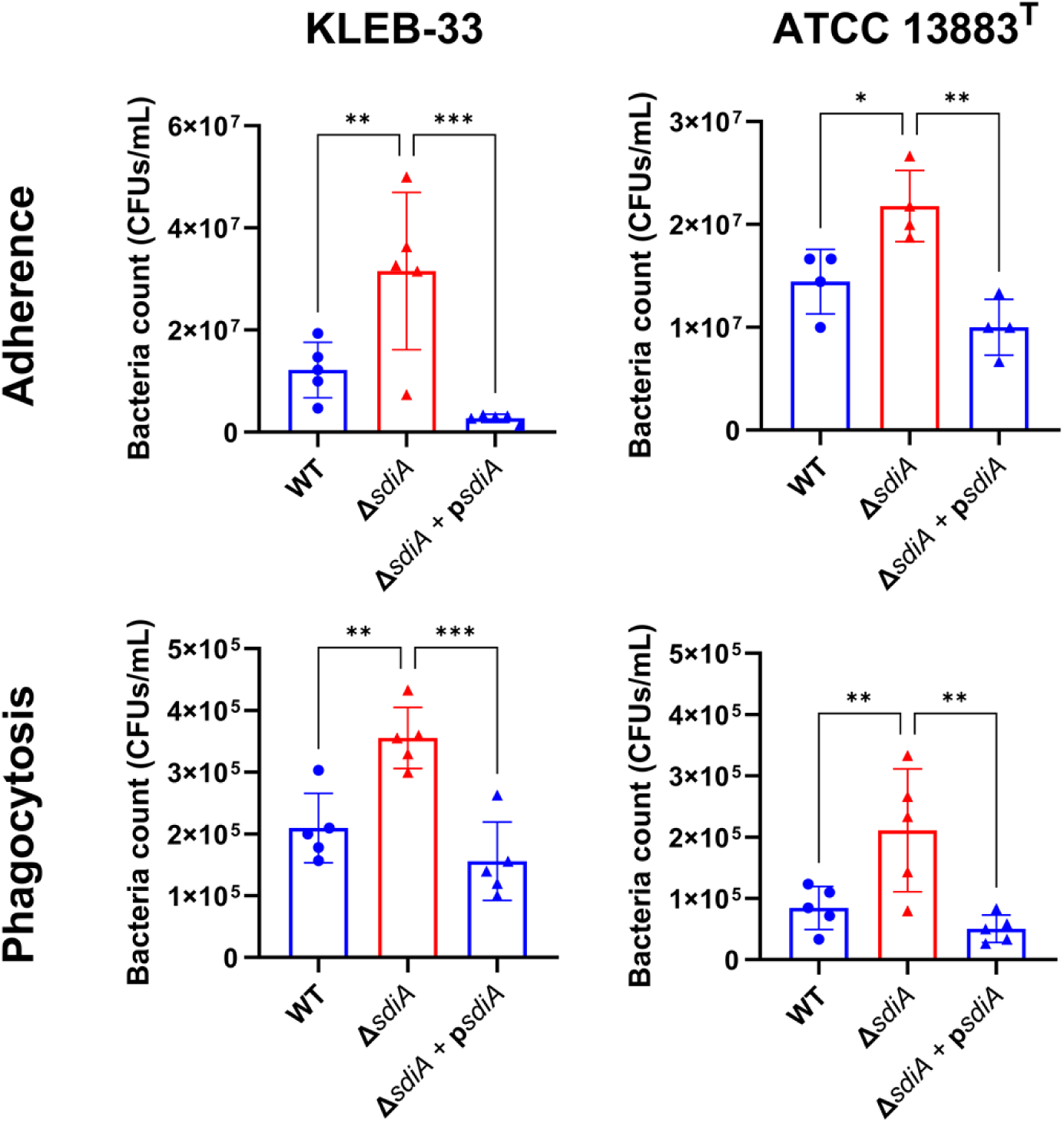
Adhesion and phagocytosis of *K. pneumoniae* WT, Δ*sdiA*, and complemented Δ*sdiA* (Δ*sdiA* + p*sdiA*) strains infecting murine iBMDMs, quantified as Colony Forming Units per mL (CFUs/mL) recovered from macrophage plates. Left panels correspond to KLEB-33 and right panels to ATCC 13883^T^. From top to bottom: it is represented the adhesion (30 min post-infection) and phagocytosis (60 min post-infection). Experiment was performed in triplicate with N=2.

### 4.2. Δ*sdiA* shows lower uronic acid quantities

Capsular polysaccharides (CPS) are key virulence factors that protect bacteria from phagocytic uptake, and reduced capsule levels are associated with increased adhesion and surface colonisation (3,7). Accordingly, alterations in capsule production represent a plausible mechanism underlying the differences in phagocytosis and adhesion observed between WT and Δ*sdiA* strains.

To investigate this, CPS levels were estimated by uronic acid quantification using the carbazole method, a widely used approach for CPS quantification based on the presence of uronic acid residues such as glucuronic or galacturonic acid in most capsule types (39). Although over 80 capsular types have been described, some exceptions exist; for instance, type K57 has been reported to lack or contain minimal uronic acid (39). However, literature indicates that K51 (KLEB-33 strain) and K3 (ATCC 13883^T^ strain) do contain uronic acid (40,41), supporting the applicability of this assay to the strains used in this study.

Uronic acid quantification revealed a significant decrease in Δ*sdiA* strains in both genetic backgrounds compared to their respective WT controls (**Figure 3**). This reduction was consistently observed in both KLEB-33 and ATCC 13883T strains, indicating that the effect is independent of strain-specific traits. The magnitude of the decrease was comparable between backgrounds, suggesting a robust and reproducible phenotype associated with *sdiA* deletion. In contrast, plasmid-based complementation (Δ*sdiA* + p*sdiA*) restored uronic acid levels to those observed in WT strains, confirming that the reduction in CPS content is specifically linked to the absence of SdiA. Overall, these data demonstrate that deletion of *sdiA* leads to a consistent reduction in CPS-associated uronic acid content across different *K. pneumoniae* backgrounds, which may contribute to the increased adhesion and phagocytosis observed in Δ*sdiA* strains.

**Figure 3.**
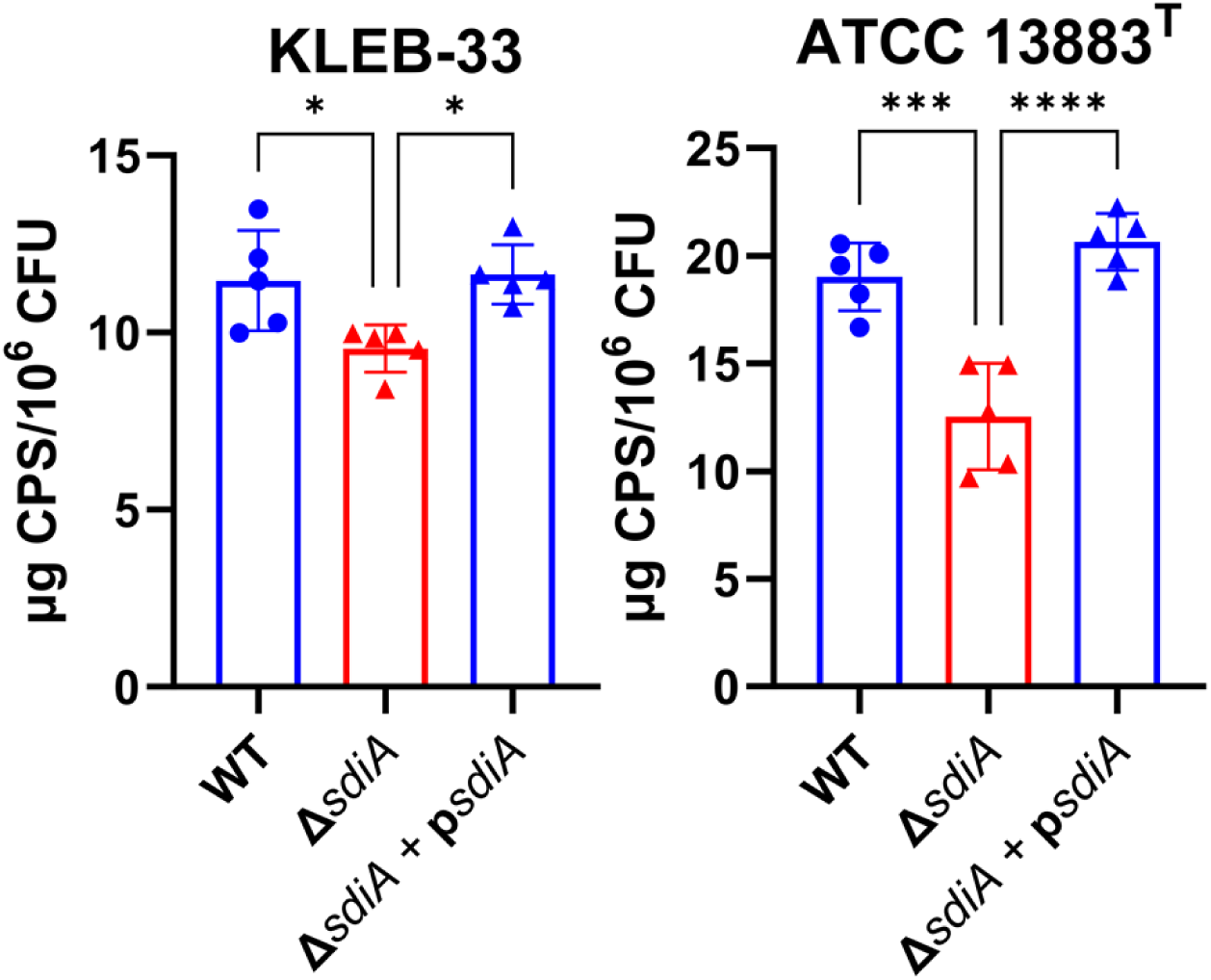
Uronic acid quantification of *K. pneumoniae* KLEB-33 and ATCC 13883^T^ WT, Δ*sdiA*, and complemented Δ*sdiA* (Δ*sdiA* + p*sdiA*) strains. Uronic acid content was determined using the carbazole method by measuring Abs_530 nm_ and expressed as µg of Capsular Polysaccharides per 10^6^ CFU (µg CPS/10^6^ CFU). Experiments were performed twice with N=3.

### 4.3. Δ*sdiA* strains exhibit enhanced intracellular survival and replication in macrophages

Given the increased adhesion and phagocytosis observed in Δ*sdiA* strains (**Figure 1** and **Figure 2**), we next investigated whether these differences translated into altered intracellular survival within macrophages. Intracellular survival was assessed in iBMDMs infected at MOI 50 and analysed at 60, 180 and 300 min post-infection by CFU enumeration.

Δ*sdiA* strains showed significantly increased intracellular survival at 180- and 300-min post-infection in both genetic backgrounds (**Figure 4A**). Importantly, this increase is not solely attributable to higher initial uptake, as intracellular CFU counts were quantified starting from 60 min post-infection, after removal of extracellular bacteria. At this time point, both WT and Δ*sdiA* populations were already intracellular, allowing direct assessment of intracellular bacterial dynamics. CFU counts increased from 60 to 180 min in both strains, indicating active intracellular proliferation rather than simple persistence. This was followed by a decrease at 300 min, suggesting either bacterial killing or loss of host cell integrity. Notably, Δ*sdiA* strains exhibited a significantly steeper increase between 60 and 180 min, resulting in higher intracellular bacterial loads compared to WT. This differential slope indicates enhanced intracellular replication capacity of the mutant, in addition to the higher initial bacterial uptake observed for this strain.

**Figure 4.**
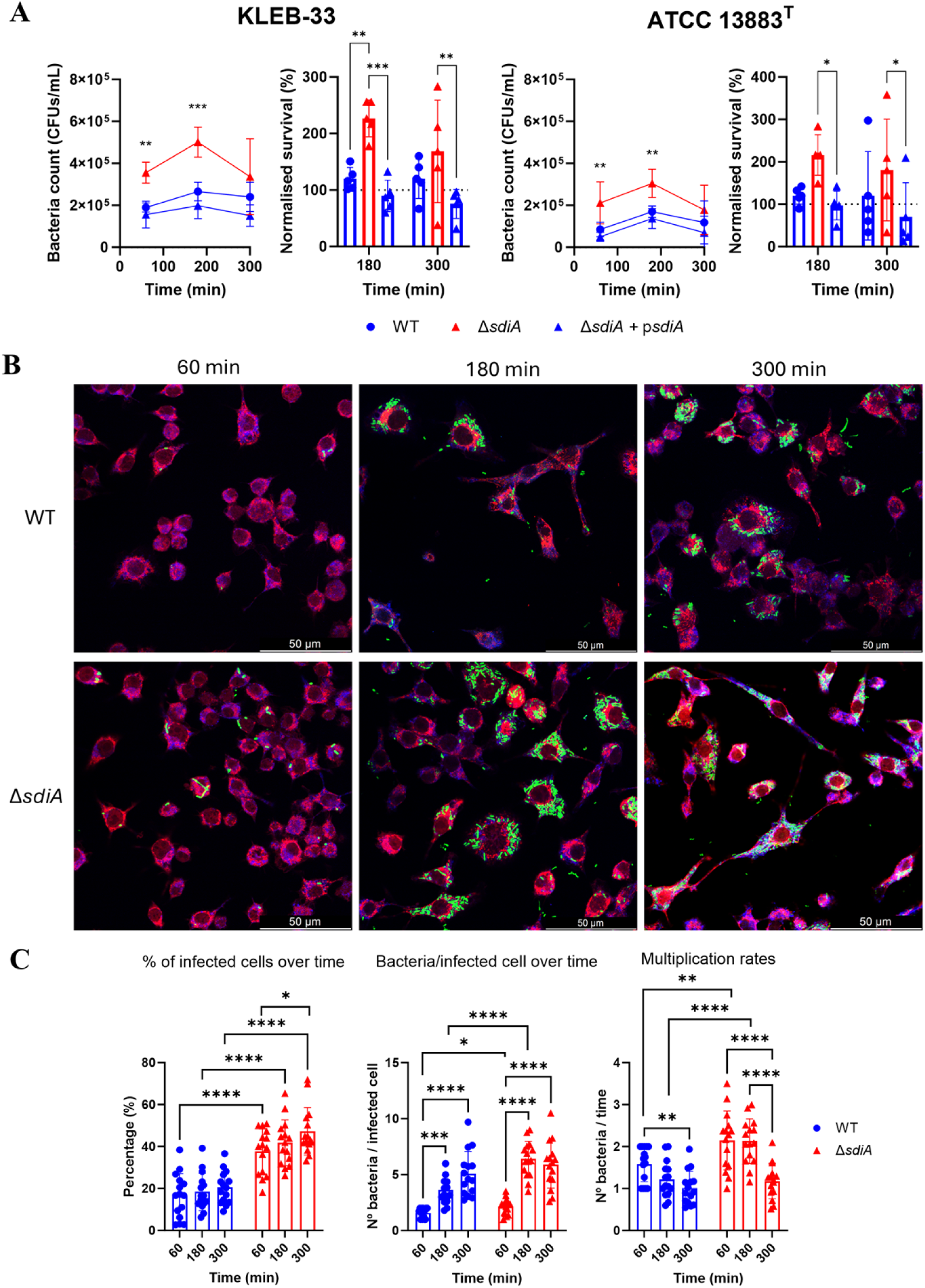
iBMDMs infected with *K. pneumoniae* WT, Δ*sdiA*, and complemented Δ*sdiA* (Δ*sdiA* + p*sdiA*) strains at MOI 50 and analysed at for 60, 180 and 300 min post-infection in the presence of gentamicin (100 µg/mL). (**A**) Intracellular survival dynamics and relative intracellular survival at 180 and 300-min, expressed as percentage relative to WT (set to 100 %, dotted line), for KLEB-33 and ATCC 13883^T^. (**B**) Representative confocal images of KLEB-33 infecting iBMDMs. Bacteria are shown in green (GFP), lysosomes in blue (Lamp1, AlexaFluor405), and mitochondria in red (MitoTracker®). Two samples per condition were analysed, with 5 fields/sample acquired at 1000x magnification. (**C**) Quantification of infection parameters: percentage of infected cells (left), number of bacteria per infected cell (middle), and bacterial multiplication rate (right), calculated as the number of intracellular bacteria per cell divided by incubation time. Image analysis was performed using Leica Application Suite Office (v1.4.6.28433) and Fiji (v1.54p) software. Only bacteria colocalising with Lamp1 was considered intracellular.

To further dissect these dynamics, confocal microscopy was used to monitor infection at the single-cell level (**Figure 4B-C**). Δ*sdiA* strains infected a significantly higher proportion of macrophages (approximately 20 % increase) compared to WT across time points. Moreover, quantification revealed that the number of bacteria per infected cell increased markedly from 60 to 180 min in both WT and mutant strains, consistent with intracellular proliferation. However, Δ*sdiA* strains consistently exhibited higher bacterial loads per infected cell, indicating that the increased CFU counts reflect not only a greater number of infected cells but also a higher intracellular burden per cell. Importantly, calculation of multiplication rates confirmed that Δ*sdiA* bacteria replicate more efficiently within macrophages, particularly at early time points (60-180 min), before decreasing at 300 min (**Figure 4C**).

### 4.4. Enhanced intracellular burden of Δ*sdiA* correlates with increased macrophage death

In the intracellular survival experiments, a decrease in CFU counts was observed between the 180- and 300-min time points (**Figure 4A**) across all strains. This reduction could reflect either bacterial killing by macrophages or macrophage death leading to bacterial release into the gentamicin-containing supernatant, where bacteria would subsequently be eliminated. As a result, CFU-based measurements alone do not allow discrimination between bacterial clearance and host cell death. To address this limitation, macrophage viability was assessed at the end of infection using the Neutral Red assay (16). Neutral Red enters cells by passive diffusion and accumulates in lysosomes, where it becomes protonated and retained due to ATP-dependent pH gradients. Loss of membrane integrity or metabolic activity results in reduced dye retention. Therefore, the amount of retained Neutral Red is proportional to the number of viable macrophages.

Quantification of cell viability revealed a clear reduction in macrophage survival upon infection with Δ*sdiA* strains compared to WT and complemented strains in both KLEB-33 and ATCC 13883T backgrounds (**Figure 5**). While all infected conditions showed decreased viability relative to uninfected controls, Δ*sdiA*-infected macrophages consistently exhibited the lowest viability levels. This trend was reproducible across independent experiments and was observed in both strain backgrounds, indicating a robust phenotype. Importantly, the reduced viability observed in Δ*sdiA* infections correlates with the higher intracellular bacterial burden previously detected (**Figure 4A**). Taken together, these data support the interpretation that the decrease in CFU counts at later time points could be associated with macrophage death and subsequent bacterial release into the antibiotic-containing supernatant.

**Figure 5.**
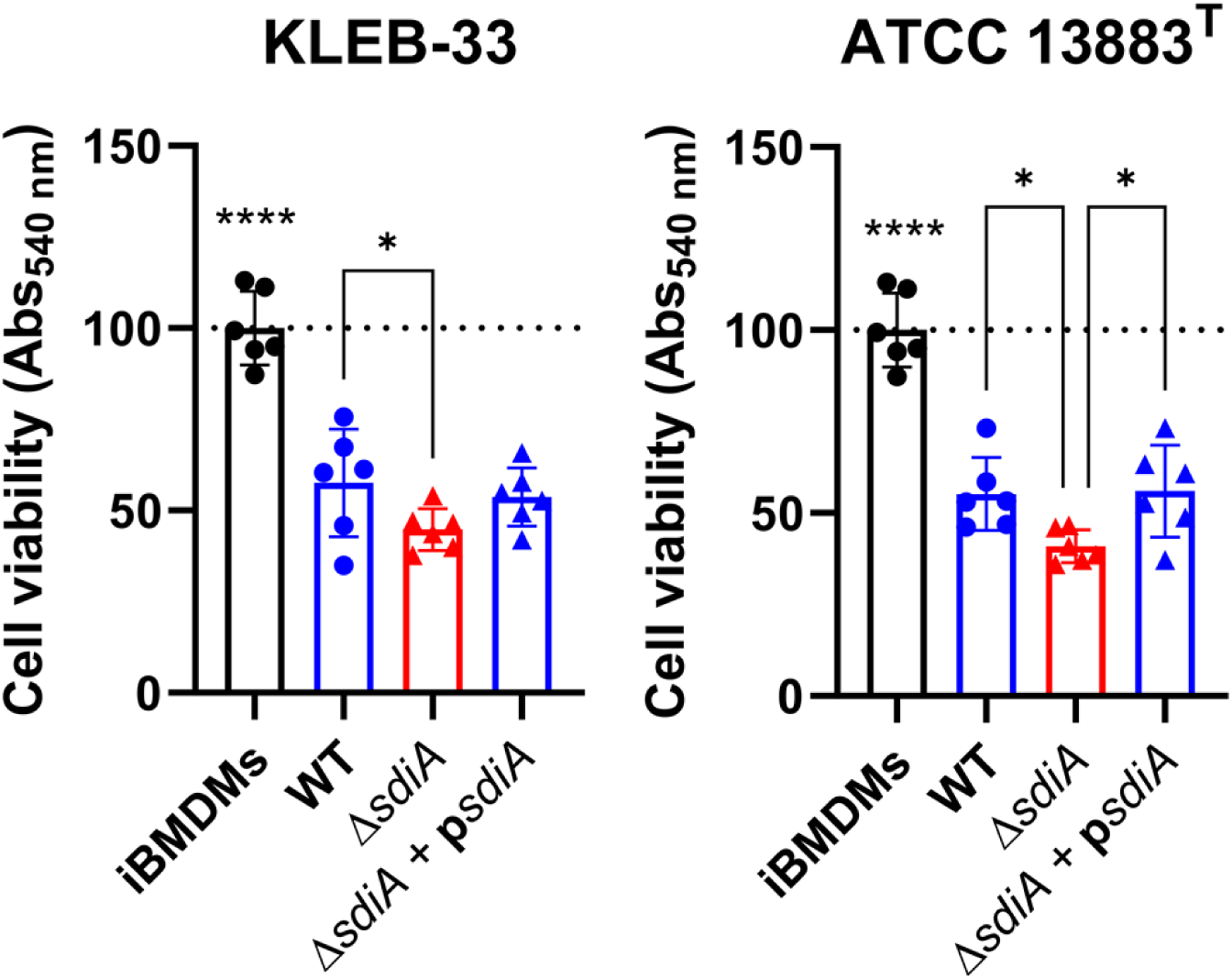
Viability of iBMDMs after 300 min of infection with *K. pneumoniae* strains. Macrophages infected with KLEB-33 (left) or ATCC 13883^T^ (right) WT, Δ*sdiA*, and complemented Δ*sdiA* (Δ*sdiA* + p*sdiA*) strains were assessed using the Neutral Red assay. Cells were incubated with Neutral Red (40 µg/mL) for 2 h, and dye retention was quantified spectrophotometrically at Abs_540 nm_ dissolving the colorant and expressed as percentage relative to uninfected controls (set to 100 %, dotted line). Experiments were performed in triplicate with N=2.

### 4.5. Δ*sdiA* does not predate *E. coli* compared to WT strains

Our previous results suggested that SdiA may influence the expression of virulence-associated factors involved in intracellular survival. To further explore whether SdiA affects bacterial competitive behaviour and the development of such factors, we performed interbacterial competition assays using a susceptible prey strain (15). Notably, several systems implicated in interbacterial interactions, such as T6SS, siderophore production, and bacteriocins, also contribute to interactions with eukaryotic hosts, including within phagosomal environments (6,15,42,43).

Killing assays were carried out using *K. pneumoniae* WT, Δ*sdiA* and complemented strains against an *E. coli* MG1655 prey strain. In both KLEB-33 and ATCC 13883T backgrounds, WT strains caused a clear reduction in prey recovery compared to the negative control, corresponding to an approximate decrease of half a log in CFU counts (**Figure 6**). A similar reduction was observed for the complemented strains, indicating restoration of the WT phenotype. In contrast, Δ*sdiA* strains showed no detectable killing activity, as prey CFU counts were comparable to those of the negative control in both strain backgrounds. This lack of reduction in prey survival was consistent across independent experiments and indicates a loss of interbacterial competitive capacity in the absence of SdiA. Overall, these results demonstrate that deletion of *sdiA* abolishes the ability of *K. pneumoniae* to reduce *E. coli* viability under the conditions tested.

**Figure 6.**
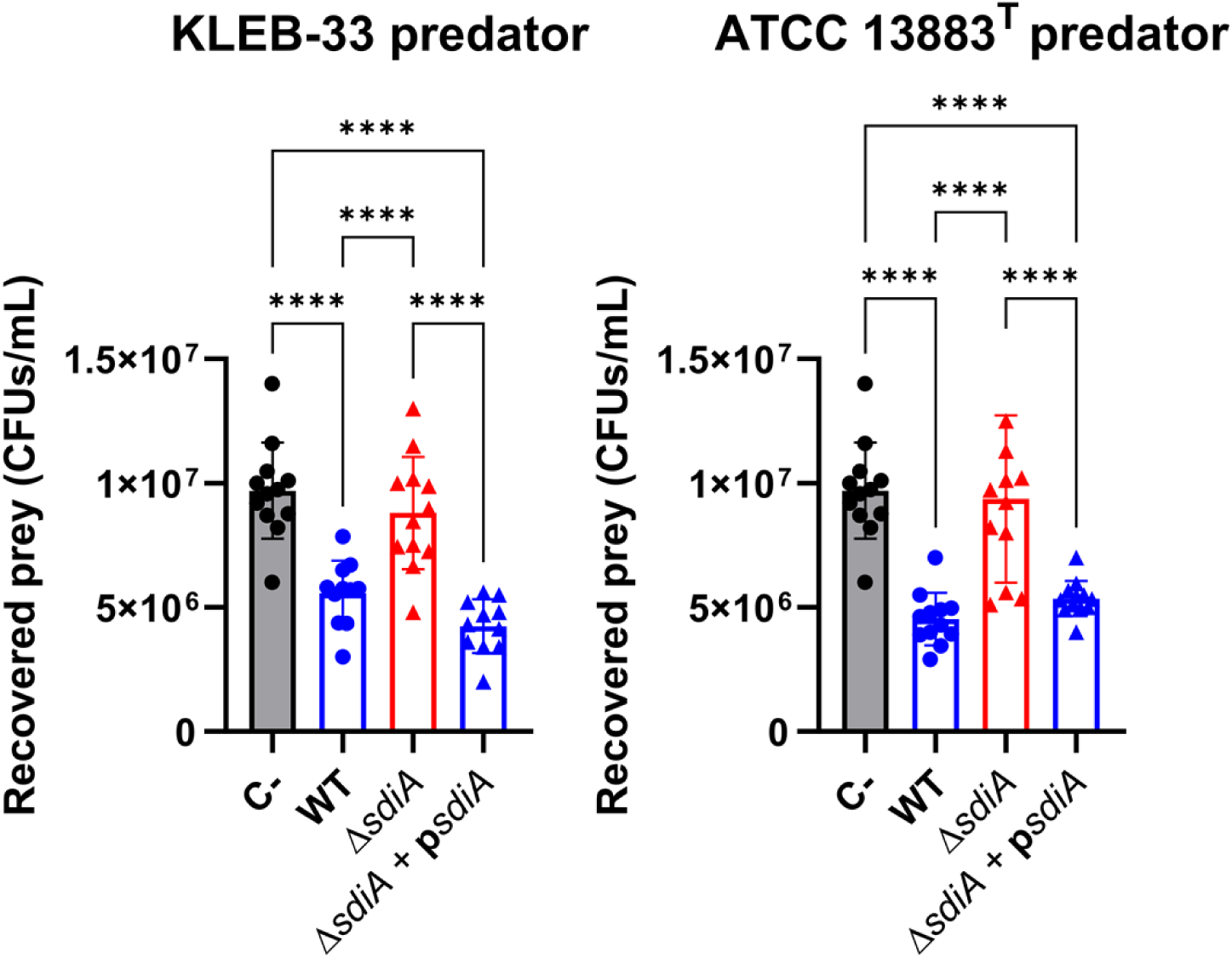
Interbacterial competition assays using *E. coli* MG1665 Strep^R^ as prey and *K. pneumoniae* WT, *ΔsdiA*, and complemented Δ*sdiA* (Δ*sdiA* + p*sdiA*) strains as predators. Predator and prey were mixed at a 10:1 ratio and incubated on agar plates (1.5 %) without nutrients for 5 h. Prey survival was quantified as CFU/mL after recovery on LB agar supplemented with streptomycin (100 µg/mL). The negative control (C-) corresponds to prey incubated alone under identical conditions. Susceptibility of predator strains to streptomycin was verified prior to the assay. Experiments were performed 4 times with N=3.

### 4.6. iBMDMs defence mechanisms are not impaired during Δ*sdiA* infection

Although interbacterial competition assays are frequently used as a proxy to assess the activity of bacterial offensive systems, they do not fully recapitulate the conditions encountered during macrophage infection. Our results indicate that the increased adhesion and intracellular survival observed in Δ*sdiA* strains are not associated with enhanced interbacterial killing capacity, suggesting that their improved persistence within macrophages is unlikely to be driven by classical antibacterial offensive systems. Instead, SdiA may influence the ability of *K. pneumoniae* to modulate or evade macrophage defence mechanisms once inside the host cell. To explore this possibility, we focused on two major antimicrobial responses: ROS production and cytokine secretion (6,13). ROS provides rapid, non-specific antimicrobial activity, whereas cytokines primarily reflect and regulate macrophage activation states, and *in vivo* contribute to the coordination of systemic immune responses and recruitment of additional immune effector cells.

Regarding ROS production, we quantified both total intracellular ROS and mitochondrial ROS at 300 min post-infection using H₂DCF-DA and MitoSOX, respectively. These complementary approaches allow discrimination between global oxidative responses and mitochondria-specific ROS production, which is a major contributor to antimicrobial activity during macrophage infection. In both KLEB-33 and ATCC 13883^T^ backgrounds, infection with *K. pneumoniae* led to a clear increase in ROS levels compared to uninfected iBMDMs, confirming activation of macrophage antimicrobial responses (**Figure 7**). Importantly, Δ*sdiA*-infected macrophages consistently exhibited significantly higher levels of both total and mitochondrial ROS than those infected with WT strains. This trend was observed across both strain backgrounds and for both ROS measurements, indicating a robust and reproducible phenotype. Complementation of *sdiA* restored ROS levels to those observed in WT infections, confirming that the effect is specifically associated with the absence of SdiA (**Figure 7**). The increased ROS production in Δ*sdiA* infections is consistent with confocal microscopy data showing a higher intracellular bacterial burden in these conditions (**Figure 4**), suggesting that macrophages mount a stronger oxidative response proportional to bacterial load. Notably, mitochondrial ROS followed the same pattern as total ROS, indicating that mitochondrial activation contributes substantially to the total oxidative response during infection.

**Figure 7.**
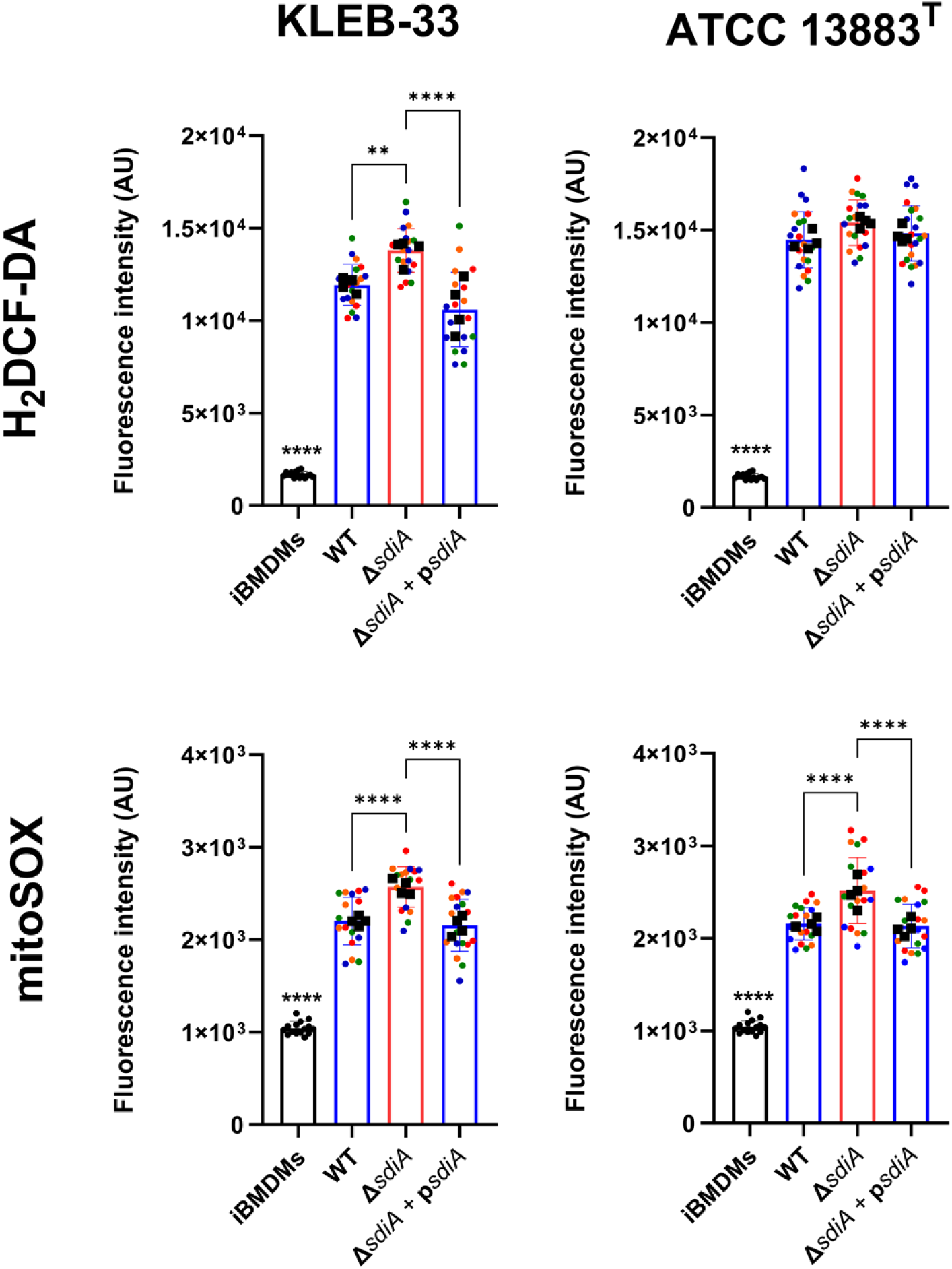
ROS production at 300 min post-infection in iBMDMs infected with *K. pneumoniae* KLEB-33 (left) and ATCC 13883^T^ (right) WT, Δ*sdiA*, and complemented Δ*sdiA* (Δ*sdiA* + p*sdiA*) strains. Total intracellular ROS (top) was measured using 2’,7’-dichlorodihydrofluorescein diacetate (H_2_DCF-DA), which fluoresces upon oxidation. Mitochondrial ROS (bottom) was quantified using MitoSOX reagent. Fluorescence intensity is expressed as Arbitrary Units (AU). Uninfected iBMDMs controls showed significant differences compared to all infected conditions. Data are represented as superplots: each colour represents an independent experiment, and black squares indicate experiment means. Experiments were performed four times with N=4, and reproducibility was confirmed by CFU quantification.

As a complementary readout of host defence activation, we analysed key pro-inflammatory mediators (mCXCL10, mTNFα and mIL-1β) in supernatants from infected iBMDMs at 300 min post-infection. These cytokines were selected as representative markers of distinct but complementary inflammatory pathways: mTNFα as a central mediator of acute inflammation, mCXCL10 as a chemokine associated with interferon-related responses and immune cell recruitment, and mIL-1β as an inflammasome-dependent cytokine (44,45). Cytokine levels were measured by ELISA, allowing quantitative comparison across infection conditions.

In both KLEB-33 and ATCC 13883^T^ backgrounds, infection with *K. pneumoniae* induced robust cytokine production compared to uninfected controls (**Figure 8A**). Importantly, macrophages infected with Δ*sdiA* strains produced significantly higher levels of mCXCL10 and mTNFα than those infected with WT strains, potentially as a consequence of its increased intracellular load. This increase was consistent across both strain backgrounds, indicating a reproducible enhancement of the pro-inflammatory response. In contrast, no significant differences were observed in mIL-1β production between WT and Δ*sdiA* infections. To further dissect the underlying signalling pathways, THP1-Dual reporter cells were used to assess NF-κB and IFNAR pathway activation (**Figure 8B**), as these pathways are key regulators of the pro-inflammatory and interferon-associated responses reflected by the cytokines measured. Consistent with the cytokine data, NF-κB signalling was significantly increased in Δ*sdiA* infections compared to WT. In contrast, IFNAR signalling remained unchanged, indicating that type I interferon responses were not enhanced in the absence of SdiA.

**Figure 8.**
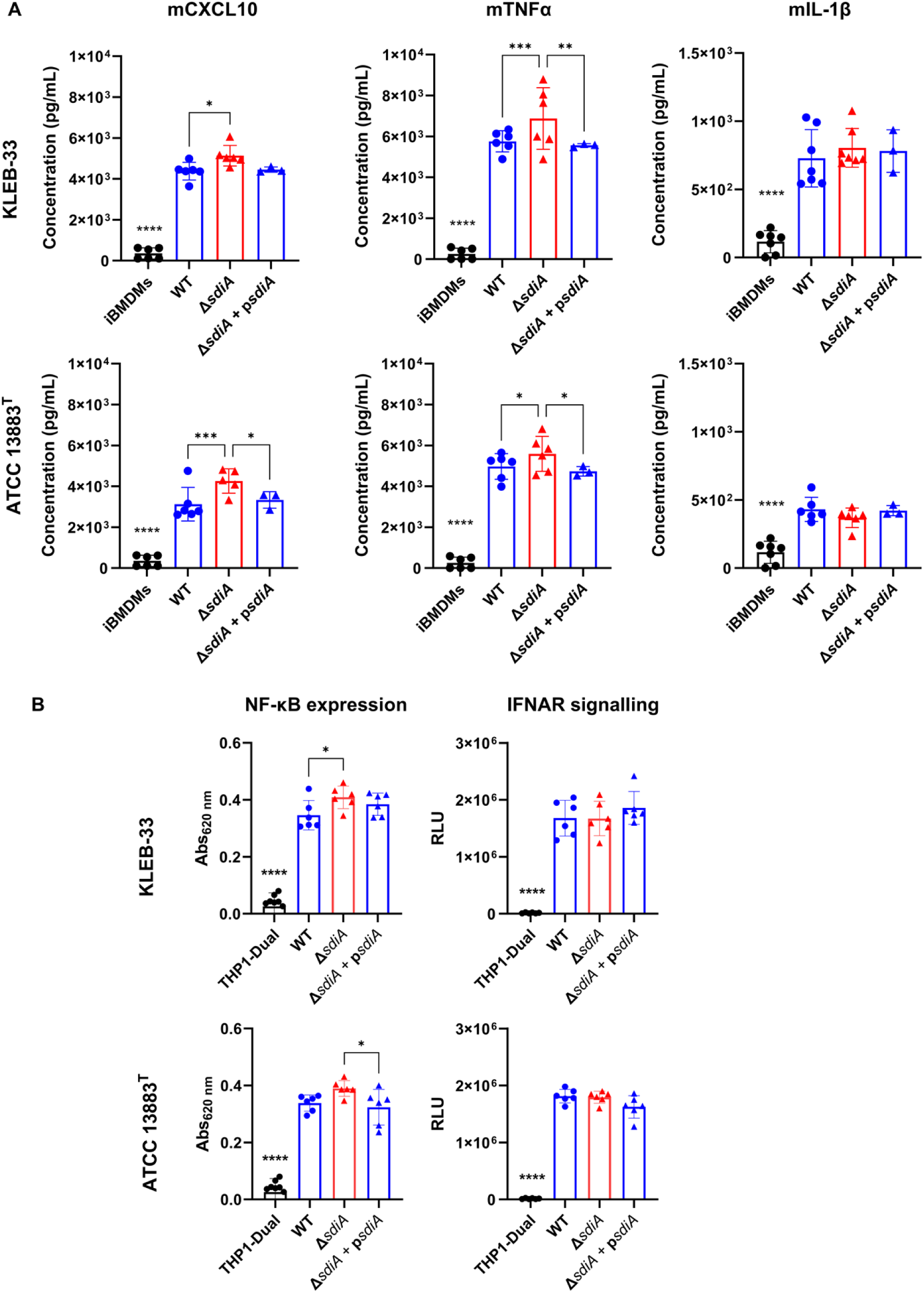
Cytokine production in iBMDMs (**A**) and signalling pathway activation in THP1-Dual reporter cells (**B**) at 300 min post-infection with *K. pneumoniae* KLEB-33 (top) and ATCC 13883^T^ (bottom) WT, Δ*sdiA*, and complemented Δ*sdiA* (Δ*sdiA* + p*sdiA*) strains. (**A**) ELISA quantification of mCXCL10, mTNFα, and mIL-1β levels in in cell culture supernatants. (**B**) NF-κB activation (left) and IFNAR signalling (right), measured as Relative Light Units (RLU). Uninfected iBMDMs controls showed significant differences compared to all infected conditions. Experiments were performed in triplicate with N=2, and reproducibility was confirmed by CFU quantification.

## 5. Discussion

In this study, we investigated the role of the orphan quorum-sensing receptor SdiA in host-pathogen interactions, which remain poorly characterised in *K. pneumoniae*. Our findings demonstrate that SdiA plays a key role in modulating bacterial interactions with phagocytic cells, influencing phagocytosis, intracellular survival, and host cell responses.

We observed that deletion of *sdiA* significantly increased susceptibility to phagocytosis in both macrophage and amoeba models, supporting a conserved role of SdiA in limiting phagocytic uptake across distinct host systems. These results suggest that SdiA contributes to resistance against phagocytic predation, thereby promoting bacterial survival against innate immune-like clearance mechanisms. This phenotype is consistent with previous reports describing increased adhesion in SdiA-deficient strains across multiple species. For instance, increased adhesion has been associated with elevated type-1 fimbriae expression in *K. pneumoniae* (9). Similarly, *Enterobacter cloacae sdiA* mutants adhered more strongly to rice roots (25,46); and in *E. coli*, Δ*sdiA* strains showed increased adhesion to HEp-2 epithelial cells (47). In contrast, another study reported reduced adhesion to HeLa cells in an SdiA-deficient *S. enterica* strain, highlighting potential species-specific regulatory effects (11).

Given the central role of the polysaccharide capsule in preventing phagocytosis (3), we assessed capsule production through uronic acid quantification. Δ*sdiA* strains consistently displayed reduced uronic acid levels, indicating decreased capsule production. This phenotype provides a plausible mechanistic explanation for the increased adhesion and phagocytosis observed in amoeba and macrophage models. These findings are in agreement with previous studies showing reduced resistance of Δ*sdiA* strains to serum complement-mediated killing, consistent with reduced capsule production (5). Moreover, a similar role for SdiA in capsule regulation was reported in *E. coli* (10), where Δ*sdiA* strains showed reduced floatability in Percoll gradients, indicative of decreased capsule production. However, contrasting results have been reported in *Cronobacter sakazakii*, where SdiA deficiency increased CPS biosynthesis gene expression (48), suggesting that SdiA-dependent regulation of capsule biosynthesis may be context- and species-dependent.

Following phagocytosis, we investigated bacterial intracellular behaviour within macrophages. Δ*sdiA* strains exhibited enhanced intracellular survival and replication, resulting in increased bacterial burden and higher levels of macrophage death. The enhanced intracellular survival of Δ*sdiA* strains could arise from a combination of an increased initial phagocytic uptake, higher bacterial load per cell, and elevated intracellular replication rates. Rather than indicating a classical increase in virulence, these results point to enhanced intracellular fitness of the mutant strain within the macrophage environment. This higher intracellular persistence contrasts with the behaviour of Δ*sdiA* strains in interbacterial competition assays, where the mutant displayed a complete loss of killing activity against *E. coli*. Comparable findings have been reported in mixed-species biofilms, where *E. coli* strains lacking *sdiA* exhibited reduced competitive capacity against *Pseudomonas aeruginosa* compared to WT strains (10). Similarly, reduced virulence has been described for Δ*sdiA* strains in other infection models, including complete loss of mouse mortality in *S. enterica* (11), as well as depletion of *Galleria mellonella* killing in *K. pneumoniae*, although this effect was strain- and condition-dependent (5).

Together, these observations reveal a dissociation between interbacterial competitiveness and intracellular fitness, suggesting that SdiA regulates distinct sets of functions depending on the biological context. While SdiA appears to promote traits associated with bacterial competition, its absence enhances traits that favour intracellular persistence. Despite the loss of interbacterial killing capacity, macrophages infected with Δ*sdiA* strains mounted an enhanced oxidative response, with increased production of both total and mitochondrial reactive oxygen species (ROS). This indicates that the enhanced intracellular survival of the mutant is not due to suppression of host antimicrobial responses, despite the known ability of *K. pneumoniae* to counteract ROS within the KCV through mechanisms such as T6SS-mediated disruption of mitochondrial networks and the expression of ROS-detoxifying enzymes (8,14,24). One possible explanation is that Δ*sdiA* bacteria may better tolerate oxidative stress once generated; however, this remains speculative and is not supported by current evidence from *E. coli*, where SdiA has been associated with increased expression of ROS defence systems (8,24).

In addition to ROS, Δ*sdiA* infections elicited enhanced pro-inflammatory responses, characterised by increased NF-κB activation, which can lead to the higher secretion of mTNFα and mCXCL10 reported (45). In contrast, no significant differences were observed in mIL-1β production. Activation of the inflammasome and subsequent release of mature mIL-1β typically require a secondary signal, such as cytosolic damage or vacuolar disruption induced by bacterial effectors (44). Therefore, these results suggest that Δ*sdiA* strains, despite inducing stronger NF-κB activation, may lack the capacity to induce higher cytosolic damage compared to WT strains. Consistent with this interpretation, the reduced interbacterial killing capacity of the mutant may reflect a diminished ability to induce intracellular damage signals.

Overall, our findings indicate that enhanced intracellular survival of Δ*sdiA* strains is not driven by suppression of host defence mechanisms, but rather by increased bacterial uptake and replication, which may overwhelm macrophage bactericidal capacity. These results highlight a complex and context-dependent role of SdiA in *K. pneumoniae* pathogenesis, where the regulation of virulence-associated traits differs between interbacterial and host-associated environments. Further studies will be required to elucidate the molecular mechanisms underlying SdiA-dependent regulation and to determine how these pathways contribute to infection dynamics *in vivo*.

## 6. Conclusions

This study identifies SdiA as a negative regulator of phagocytosis and intracellular survival in *K. pneumoniae*, likely acting in part through the positive regulation of capsular polysaccharide (CPS) production, among other possible mechanisms. Deletion of *sdiA* resulted in reduced uronic acid content, indicative on diminished capsule production, consistent with increased susceptibility to amoeba predation as well as enhanced adhesion to and phagocytic uptake by murine macrophages.

Despite this increased uptake, *sdiA*-deficient strains exhibited enhanced intracellular replication and induced higher levels of macrophage death. Notably, these effects were not associated with suppression of host antimicrobial responses, as mutant-infected macrophages displayed increased ROS production and elevated levels of pro-inflammatory cytokines. In parallel, Δ*sdiA* strains showed reduced interbacterial competitive capacity against *E. coli*, indicating that their enhanced intracellular survival is unlikely to be driven by classical antibacterial offensive mechanisms.

Together, these results indicate that the increased intracellular fitness of Δ*sdiA* strains arises from a combination of enhanced uptake and intracellular proliferation, which may overwhelm macrophage bactericidal capacity rather than actively suppressing host defence mechanisms. More broadly, our findings highlight a context-dependent role of SdiA in *K. pneumoniae* virulence, revealing a dissociation between interbacterial competitiveness and intracellular survival. This work underscores the importance of host-pathogen context when interpreting bacterial virulence traits and refines our understanding of how *K. pneumoniae* adapts to the intracellular environment.

## Author Contributions

**Sergio Silva-Bea**: Conceptualization, Data curation, Formal analysis, Investigation, Methodology, Resources, Visualization, Writing - original draft. **Ana Otero** and **Manuel Romero**: Conceptualization, Funding acquisition, Project administration, Resources, Supervision, Validation, Writing - review & editing. **Ricardo Calderon-Gonzalez** and **Joana Sa-Pessoa**: Data curation, Formal analysis, Investigation, Methodology, Resources**. Jose A. Bengoechea**: Data curation, Methodology, Resources Conceptualization, Funding acquisition, Validation, Project administration.

## Funding

This research was funded by grants from Conselleria de Cultura, Educación, e Ordenación Universitaria, Xunta de Galicia, (ED431B2023/4). Sergio Silva-Bea was supported with the “Formación de Profesorado Universitario” program (FPU21/01147), the “Ayudas de movilidad para estancias breves” (EST24/00156) and “Convocatoria de estadías e axudas para asistencia a congresos iARCUS 2025” grants. Jose A. Bengoechea supported this research with the projects BB/T001976/1 and BB/V007939/1 from UK Biotechnology and Biological Sciences Research Council (BBSRC).

## Data Availability Statement

The data that support the findings of this study are available from the corresponding authors upon reasonable request.

## Conflict of Interest

The authors declare that the research was conducted in the absence of any commercial or financial relationships that could be construed as a potential conflict of interest.

## Permission to reuse and Copyright

All authors have approved the manuscript before submission to publishing the figures and tables under the normative of the journal.

## Declaration of generative AI use

During the preparation of this work the authors used Google Gemini (Version 3.1 Pro, https://gemini.google.com/) for language editing purposes. All content was subsequently reviewed and approved by the authors, who take full responsibility for the manuscript.

## Notes

### Competing Interest Statement

The authors have declared no competing interest.

## References

1. Tacconelli E, Carrara E, Savoldi A, Harbarth S, Mendelson M, Monnet DL, et al. Discovery, research, and development of new antibiotics: the WHO priority list of antibiotic-resistant bacteria and tuberculosis. Lancet Infect Dis. 2018 Mar;18(3):318–27. doi:10.1016/S1473-3099(17)30753-3

2. WHO. WHO Bacterial Priority Pathogens List 2024 Bacterial Pathogens of Public Health Importance, to Guide Research, Development, and Strategies to Prevent and Control Antimicrobial Resistance. Geneva: World Health Organization; 2024.

3. González-Ferrer S, Peñaloza HF, Budnick JA, Bain WG, Nordstrom HR, Lee JS, et al. Finding Order in the Chaos: Outstanding Questions in *Klebsiella pneumoniae* Pathogenesis. Ottemann KM, editor. Infect Immun. 2021 Mar 17;89(4):e00693–20. doi:10.1128/IAI.00693-20

4. Russo TA, Marr CM. Hypervirulent *Klebsiella pneumoniae*. Clin Microbiol Rev. 2019;32(3). 10.1128/CMR.00001-19.

5. Silva-Bea S, Maseda P, Otero A, Romero M. Regulatory effects on virulence and phage susceptibility revealed by *sdiA* mutation in *Klebsiella pneumoniae*. Front Cell Infect Microbiol. 2025 Mar 13;15:1562402. doi:10.3389/fcimb.2025.1562402

6. Bengoechea JA, Sa Pessoa J. *Klebsiella pneumoniae* infection biology: living to counteract host defences. FEMS Microbiol Rev. 2019 Mar 1;43(2):123–44. doi:10.1093/femsre/fuy043

7. Dorman MJ, Feltwell T, Goulding DA, Parkhill J, Short FL. The Capsule Regulatory Network of *Klebsiella pneumoniae* Defined by density-TraDISort. Chang YF, editor. mBio. 2018 Dec 21;9(6):e01863–18. doi:10.1128/mBio.01863-18

8. Mayer C, Borges A, Flament-Simon SC, Simões M. Quorum sensing architecture network in *Escherichia coli* virulence and pathogenesis. FEMS Microbiol Rev. 2023 Jul 5;47(4):fuad031. doi:10.1093/femsre/fuad031

9. Pacheco T, Gomes AÉI, Siqueira NMG, Assoni L, Darrieux M, Venter H, et al. SdiA, a Quorum-Sensing Regulator, Suppresses Fimbriae Expression, Biofilm Formation, and Quorum-Sensing Signaling Molecules Production in *Klebsiella pneumoniae*. Front Microbiol. 2021 Jun 21;12:597735. doi:10.3389/fmicb.2021.597735

10. Mayer C, Soukarieh F, Simões M, Flament-Simon SC, Cámara M, Romero M. AHL-based QS signalling promotes uropathogenic *Escherichia coli* settlement through the de-repression of biofilm formation by SdiA. PLOS ONE. 2025 Sep 9;20(9):e0328837. doi:10.1371/journal.pone.0328837

11. Askoura M, Almalki AJ, Lila ASA, Almansour K, Alshammari F, Khafagy ES, et al. Alteration of *Salmonella enterica* Virulence and Host Pathogenesis through Targeting sdiA by Using the CRISPR-Cas9 System. Microorganisms. 2021 Dec 11;9(12):2564. doi:10.3390/microorganisms9122564

12. Cano V, March C, Insua JL, Aguiló N, Llobet E, Moranta D, et al. *Klebsiella pneumoniae* survives within macrophages by avoiding delivery to lysosomes. Cell Microbiol. 2015 Nov;17(11):1537–60. doi:10.1111/cmi.12466

13. Li Y, Li X, Wu W, Liu P, Liu J, Jiang H, et al. Insights into the roles of macrophages in *Klebsiella pneumoniae* infections: a comprehensive review. Cell Mol Biol Lett. 2025 Mar 26;30(1):34. doi:10.1186/s11658-025-00717-7

14. Sá-Pessoa J, López-Montesino S, Przybyszewska K, Rodríguez-Escudero I, Marshall H, Ova A, et al. A trans-kingdom T6SS effector induces the fragmentation of the mitochondrial network and activates innate immune receptor NLRX1 to promote infection. Nat Commun. 2023 Feb 16;14(1):871. doi:10.1038/s41467-023-36629-3

15. Storey D, McNally A, Åstrand M, sa-Pessoa Graca Santos J, Rodriguez-Escudero I, Elmore B, et al. *Klebsiella pneumoniae* type VI secretion system-mediated microbial competition is PhoPQ controlled and reactive oxygen species dependent. Parsek MR, editor. PLOS Pathog. 2020 Mar 19;16(3):e1007969. doi:10.1371/journal.ppat.1007969

16. Feriotti C, Sá-Pessoa J, Calderón-González R, Gu L, Morris B, Sugisawa R, et al. *Klebsiella pneumoniae* hijacks the Toll-IL-1R protein SARM1 in a type I IFN-dependent manner to antagonize host immunity. Cell Rep. 2022 Aug;40(6):111167. doi:10.1016/j.celrep.2022.111167

17. Liang Z, Wang Y, Lai Y, Zhang J, Yin L, Yu X, et al. Host defense against the infection of *Klebsiella pneumoniae*: New strategy to kill the bacterium in the era of antibiotics? Front Cell Infect Microbiol. 2022 Nov 24;12:1050396. doi:10.3389/fcimb.2022.1050396

18. Tomás A, Lery L, Regueiro V, Pérez-Gutiérrez C, Martínez V, Moranta D, et al. Functional Genomic Screen Identifies *Klebsiella pneumoniae* Factors Implicated in Blocking Nuclear Factor κB (NF-κB) Signaling. J Biol Chem. 2015 Jul;290(27):16678–97. doi:10.1074/jbc.M114.621292

19. Dumigan A, Cappa O, Morris B, Sá Pessoa J, Calderon-Gonzalez R, Mills G, et al. In vivo single-cell transcriptomics reveal *Klebsiella pneumoniae* skews lung macrophages to promote infection. EMBO Mol Med. 2022 Nov 7;14(12):EMMM202216888. doi:10.15252/emmm.202216888

20. Grubwieser P, Hilbe R, Gehrer CM, Grander M, Brigo N, Hoffmann A, et al. *Klebsiella pneumoniae* manipulates human macrophages to acquire iron. Front Microbiol. 2023 Aug 11;14. doi:10.3389/fmicb.2023.1223113

21. Ahn D, Prince A. Participation of Necroptosis in the Host Response to Acute Bacterial Pneumonia. J Innate Immun. 2017 May;9(3):262–70. doi:10.1159/000455100 PubMed PMID: 28125817; PubMed Central PMCID: PMC5413418.

22. Ma X, Zhang S, Xu Z, Li H, Xiao Q, Qiu F, et al. SdiA Improves the Acid Tolerance of *E. coli* by Regulating GadW and GadY Expression. Front Microbiol. 2020 Jun 3;11:1078. doi:10.3389/fmicb.2020.01078

23. Zhou X, Meng X, Sun B. An EAL domain protein and cyclic AMP contribute to the interaction between the two quorum sensing systems in *Escherichia coli*. Cell Res. 2008 Sep;18(9):937–48. doi:10.1038/cr.2008.67

24. Barth E, Gora KV, Gebendorfer KM, Settele F, Jakob U, Winter J. Interplay of cellular cAMP levels, σ S activity and oxidative stress resistance in Escherichia coli. Microbiology. 2009 May 1;155(5):1680–9. doi:10.1099/mic.0.026021-0

25. Sabag-Daigle A, Dyszel JL, Gonzalez JF, Ali MM, Ahmer BMM. Identification of sdiA-regulated genes in a mouse commensal strain of *Enterobacter cloacae*. Front Cell Infect Microbiol. 2015;5(47). doi:10.3389/fcimb.2015.00047

26. Cheng C, Yan X, Liu B, Jiang T, Zhou Z, Guo F, et al. SdiA Enhanced the Drug Resistance of *Cronobacter sakazakii* and Suppressed Its Motility, Adhesion and Biofilm Formation. Front Microbiol. 2022 May 6;13:901912. doi:10.3389/fmicb.2022.901912

27. Pena RT, Blasco L, Ambroa A, González-Pedrajo B, Fernández-García L, López M, et al. Relationship Between Quorum Sensing and Secretion Systems. Front Microbiol. 2019 Jun 7;10:1100. doi:10.3389/fmicb.2019.01100

28. Shen Y, Gao S, Fan Q, Zuo J, Wang Y, Yi L, et al. New antibacterial targets: Regulation of quorum sensing and secretory systems in zoonotic bacteria. Microbiol Res. 2023 Sep;274:127436. doi:10.1016/j.micres.2023.127436

29. Zhang X, Liu B, Ding X, Bin P, Yang Y, Zhu G. Regulatory Mechanisms between Quorum Sensing and Virulence in *Salmonella*. Microorganisms. 2022 Nov 9;10(11):2211. doi:10.3390/microorganisms10112211

30. Steele-Mortimer O, Brumell JH, Knodler LA, Meresse S, Lopez A, Finlay BB. The invasion-associated type III secretion system of *Salmonella enterica* serovar Typhimurium is necessary for intracellular proliferation and vacuole biogenesis in epithelial cells. Cell Microbiol. 2002 Jan;4(1):43–54. doi:10.1046/j.1462-5822.2002.00170.x

31. Silva-Bea S, García-Meniño I, Rey S, Romero M, Fernández J, Hammerl JA, et al. Draft genome sequence of *Klebsiella pneumoniae* KLEB-33: a convergent biofilm hyperforming multiresistant strain belonging to the emerging ST16 lineage harboring multiple hypervirulence genes. Putonti C, editor. Microbiol Resour Announc. 2024 Apr 11;13(4):e01192–23. doi:10.1128/mra.01192-23

32. Paquet VE, Charette SJ. Amoeba-resisting bacteria found in multilamellar bodies secreted by *Dictyostelium discoideum:* social amoebae can also package bacteria. Kümmerli R, editor. FEMS Microbiol Ecol. 2016 Mar;92(3):fiw025. doi:10.1093/femsec/fiw025

33. Bartholomew TL, Kidd TJ, Sá Pessoa J, Conde Álvarez R, Bengoechea JA. 2-Hydroxylation of *Acinetobacter baumannii* Lipid A Contributes to Virulence. Bäumler AJ, editor. Infect Immun. 2019 Apr;87(4):e00066–19. doi:10.1128/IAI.00066-19

34. Campos MA, Vargas MA, Regueiro V, Llompart CM, Albertí S, Bengoechea JA. Capsule Polysaccharide Mediates Bacterial Resistance to Antimicrobial Peptides. Infect Immun. 2004 Dec;72(12):7107–14. doi:10.1128/IAI.72.12.7107-7114.2004

35. Rahn A, Whitfield C. Transcriptional organization and regulation of the *Escherichia coli* K30 group 1 capsule biosynthesis (*cps*) gene cluster. Mol Microbiol. 2003 Feb;47(4):1045–60. doi:10.1046/j.1365-2958.2003.03354.x

36. Murphy MP, Bayir H, Belousov V, Chang CJ, Davies KJA, Davies MJ, et al. Guidelines for measuring reactive oxygen species and oxidative damage in cells and in vivo. Nat Metab. 2022 Jun 27;4(6):651–62. doi:10.1038/s42255-022-00591-z

37. Adiba S, Nizak C, Van Baalen M, Denamur E, Depaulis F. From Grazing Resistance to Pathogenesis: The Coincidental Evolution of Virulence Factors. Ahmed N, editor. PLoS ONE. 2010 Aug 11;5(8):e11882. doi:10.1371/journal.pone.0011882

38. Pukatzki S, Kessin RH, Mekalanos JJ. The human pathogen *Pseudomonas aeruginosa* utilizes conserved virulence pathways to infect the social amoeba *Dictyostelium discoideum*. Proc Natl Acad Sci. 2002 Mar 5;99(5):3159–64. doi:10.1073/pnas.052704399

39. Hsu CR, Liao CH, Lin TL, Yang HR, Yang FL, Hsieh PF, et al. Identification of a capsular variant and characterization of capsular acetylation in *Klebsiella pneumoniae* PLA-associated type K57. Sci Rep. 2016 Aug 23;6(1):31946. doi:10.1038/srep31946

40. Chakraborty AK, Dąbrowski U, Geyer H, Geyer R, Stirm S. Primary structure of the *Klebsiella* serotype-51 capsular polysaccharide. Carbohydr Res. 1982 May;103(1):101–5. doi:10.1016/S0008-6215(82)80010-4

41. Pan YJ, Lin TL, Chen CT, Chen YY, Hsieh PF, Hsu CR, et al. Genetic analysis of capsular polysaccharide synthesis gene clusters in 79 capsular types of *Klebsiella* spp. Sci Rep. 2015 Oct 23;5(1):15573. doi:10.1038/srep15573

42. Ho BT, Dong TG, Mekalanos JJ. A View to a Kill: The Bacterial Type VI Secretion System. Cell Host Microbe. 2014 Jan 15;15(1):9–21. doi:10.1016/j.chom.2013.11.008 PubMed PMID: 24332978.

43. Russell AB, Peterson SB, Mougous JD. Type VI secretion system effectors: poisons with a purpose. Nat Rev Microbiol. 2014 Feb;12(2):137–48. doi:10.1038/nrmicro3185

44. Chen M ye, Ye X jia, He X hui, Ouyang D yun. The Signaling Pathways Regulating NLRP3 Inflammasome Activation. Inflammation. 2021 Aug;44(4):1229–45. doi:10.1007/s10753-021-01439-6

45. Yeruva S, Ramadori G, Raddatz D. NF-κB-dependent synergistic regulation of CXCL10 gene expression by IL-1β and IFN-γ in human intestinal epithelial cell lines. Int J Colorectal Dis. 2008 Mar;23(3):305–17. doi:10.1007/s00384-007-0396-6

46. Shankar M, Ponraj P, Illakkiam D, Rajendhran J, Gunasekaran P. Inactivation of the Transcriptional Regulator-Encoding Gene *sdiA* Enhances Rice Root Colonization and Biofilm Formation in Enterobacter cloacae GS1. J Bacteriol. 2013 Jan;195(1):39–45. doi:10.1128/JB.01236-12

47. Sharma VK, Bearson SMD, Bearson BL. Evaluation of the effects of *sdiA*, a luxR homologue, on adherence and motility of *Escherichia coli* O157 : H7. Microbiology. 2010 May 1;156(5):1303–12. doi:10.1099/mic.0.034330-0

48. Cao Y, Li L, Zhang Y, Liu F, Xiao X, Li X, et al. Evaluation of *Cronobacter sakazakii* biofilm formation after sdiA knockout in different osmotic pressure conditions. Food Res Int. 2022 Jan;151:110886. doi:10.1016/j.foodres.2021.110886

